# jULIEs: extracellular probes for recordings and stimulation in the structurally and functionally intact mouse brain

**DOI:** 10.1101/721548

**Authors:** Romeo Racz, Mihaly Kollo, Gabriella Racz, Ciprian Bulz, Tobias Ackels, Tom Warner, William Wray, Nikolai Kiskin, Chi Chen, Zhiwen Ye, Livia de Hoz, Ede Rancz, Andreas Schaefer

## Abstract

High signal-to-noise, scalable and minimally invasive recording and stimulation of the nervous system in intact animals is of fundamental importance to advance the understanding of brain function. Extracellular electrodes are among the most powerful tools capable of interfacing with large neuronal populations^1-3^. Neuronal tissue damage remains a major limiting factor in scaling electrode arrays, and has been found to correlate with electrode diameter across different electrode materials, such as microfabricated Michigan and Utah-style arrays^4^, MEMS and microsystems^5^, soft polymer or tungsten electrodes^6^ and Parylene C probes^7^. Small diameter ultramicroelectrodes (UMEs), while highly desirable, pose significant technical challenges such as reaching sufficient electrolyte-electrode coupling and limiting stray signal loss. To overcome these challenges, we have designed juxtacellular Ultra-Low Impedance Electrodes (jULIEs), a scalable technique for achieving high signal-to-noise electrical recordings as well as stimulation with UMEs. jULIEs are metal-glass composite UMEs thermally drawn to outer diameters (OD) of <25 µm, with metal core diameters (ID) of as little as 1 µm. We introduce a two-step electrochemical modification strategy that reduces UME coupling impedances by two orders of magnitude. Modifications enabled high signal-to-noise neural recordings *in vivo* through wires with micrometer scale core diameters. Histological and imaging experiments indicated that local vascular damage is minimal. Spikes reached amplitudes over 1 mV *in vivo*, indicating that recordings are possible in close proximity to intact neurons. Recording sites can be arranged in arbitrary patterns tailored to various neuroanatomical target structures and allowing parallel penetrations. jULIEs thus represent a versatile platform that allows for reliable recording and manipulation of neural activity in any areas of the functionally intact mammalian brain.

## Main

jULIE neural probes were assembled (**Fig. 1 and Supplementary Figure 1**) from composite glass-metal microwires. These were produced through a modified dieless Taylor-Ulitovsky drawing method^8,9^ (**Fig.1a**). Fibers were drawn from a gold or copper-filled Pyrex glass preform prepared under inert atmosphere and heated to melting temperatures until a glass-metal droplet formed. The glass capillary was pulled and spooled up at high speeds (up to 2km/min) on a drum. Depending on pulling force and drum speed, core diameters ranged between <1 and 10 µm with glass insulation thickness up to 15 µm. The resulting microwires were thus small (**Fig.1b**), well insulated (**Fig.1c,j**) and continuously conductive over several hundred meters, yet remained flexible with ∼500 µm bending radius (**Fig.1d**). They featured smooth exteriors (**Fig.1c**) and had well-defined glass-metal boundaries (Fig.1j). Before assembling into recording arrays (**Supplementary figure 1**) individual wires were prepared as follows. Wires were rewound from the spooling drum. Stacks of wires were then embedded in a soluble thermoplastic (**see methods**) and polished at sharp angles of approximately 30 degrees (**Fig.1c, Supplementary figure 1a,b,c**) to facilitate insertion and penetration into tissue^10,11^.

**Figure 1.**
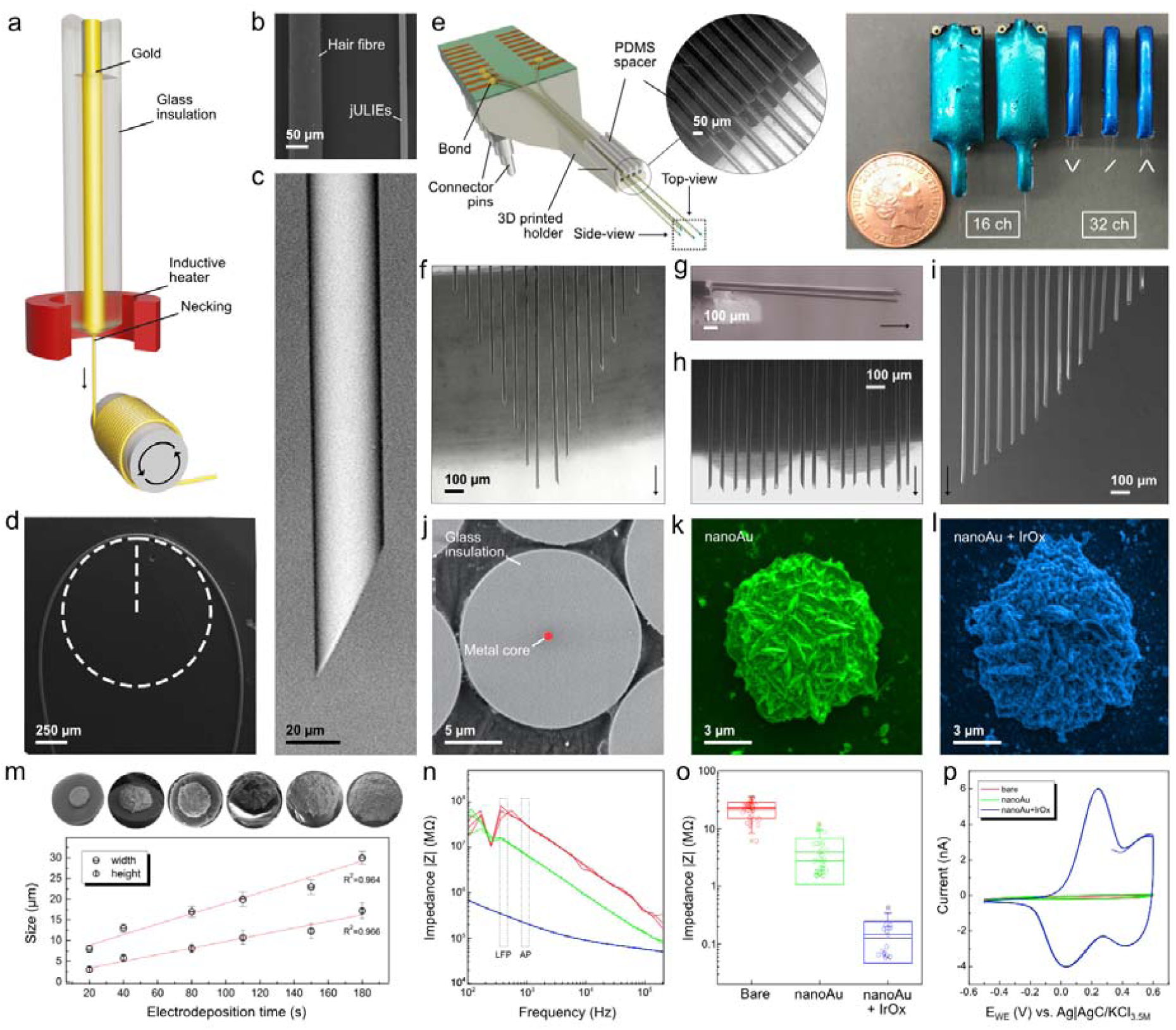
**(a)** Apparatus and principle of glass-insulated ultramicroelectrode fabrication by high-speed dieless drawing. A preform containing the metal core and glass insulation is heated and the drop formed at the necking is continuously spooled up on a drum. **(b)** SEM image comparing diameters of a human hair and a jULIE fiber. **(c)** Side-view SEM of a sharpened wire before electrochemical preparation. **(d)** SEM image of the bending radius of approx. 500 µm of a single fiber. **(e)** 3D CAD model and actual assembled jULIEs with custom layout of 16 and 32 channels and standard 128 channel probes with gold and copper cores. **(f-i)** Top-view SEM micrographs of customized recording site layouts featuring different planar tip geometries with laterally dense packing at 50 µm inter-shank distance. **(j-l)** Front-view SEM of flat polished bare fiber **(j)**, modified with nanoAu for 35 s **(k)** and nanoAu modified with IrOx **(l)**.**(m)** SEM micrograph of progressive alteration of the initial core with nanoAu by controlling electrodeposition time, nanoAu building up preferentially as a hemisphere. Electrochemical impedance spectroscopy of surface preparation stages **(j-l)**: (n) Impedance Z Bode-plot between 1 Hz and 100 kHz, **(o)** impedance at 1 kHz and (p) cyclic voltammetry response at 50 mVs^-1^ of the three preparation stages.

Given the small area of the conductive core and resulting small electrical interface with the extracellular space, impedance |Z| of the bare metal core was too high for neural recordings across frequencies from 1Hz to 100 kHz (**Fig. 1n,o**) resulting in signal attenuation and increased noise^12-14^. This is a common problem of ultra-thin electrodes. Electrode-tissue coupling can, in general, be improved through alteration of the surface by physical, chemical and electrochemical means, albeit often resulting in uncontrolled geometrical alterations and mechanical failure^15^.

Both organic and inorganic materials have been used to reduce electrode impedance for larger electrode surfaces^16^. Iridium oxide (IrOx) electrodeposition for example is a well-established surface modification for extracellular recording electrodes^17-19^. Thanks to its faradaic nature and stable reversible Ir^3+^/Ir^4+^ redox couple it also provides an excellent substrate for microstimulation^20^. Here, we modified jULIEs with an initial nanostructured gold then further added a nanostructured IrOx layer to further optimize coupling to the extracellular signal.

To maintain the cylindrical profile and small footprint of the jULIE tip, our aim was to increase the specific surface area of the metal sensor and coupling to the extracellular voltage signal while keeping geometrical size increase controlled and minimal. We therefore established a three-step microwire surface modification protocol (**Fig.1j-p, Supplementary figure 1-3**): firstly, as a result of the polishing described above (**Supplementary figure 1a-c**) the metal core was exposed in a controlled and reproducible manner (**Fig.1j**). Secondly, after de-embedding, the exposed core was modified with high-rugosity gold nanostructures (nanoAu) (**Fig.1k, Supplementary Figure 2**) by electrodeposition from an additive free cyanide electrolyte (see methods). While microwires can be produced from a wide range of metals, potential chemical incompatibility with the tissue may occur, causing toxicity or corrosion. The deposition of gold nanostructures decouples the electrode-electrolyte interface from the core metal, allowing the use of custom conductive materials for the core, largely independent of their toxicity or corrosion properties. In the following, third step, gold nanostructures were further modified with a porous film of polycrystalline IrOx (**Fig.1l, Supplementary figure 3**) from an aged IrCl_4_ electrolyte by a combined cyclic voltammetry and potentiostatic electrodeposition protocol. During the first sweeps of electrodeposition, in case of transition metal oxides, nucleation centers develop (as shown e.g. for MnO_2_^21^) and charge is being stored with each voltammetry cycle. Similarly, for IrOx (**Supplementary figure 3a**) cyclic voltammetry facilitated development of IrOx nucleation centers on nanoAu; subsequent potentiostatic pulses further accelerated buildup (**Supplementary figure 3b**).

The resulting films of polycrystalline IrOx (**Supplementary figure 3d-f**) evenly covered the nanoAu substrate (**Fig.1l**). Depending on the dimensions of the nanoAu deposit (Fig.1m) and IrOx film thickness, in combination, nanoAu+IrOx reduced interfacial impedances up to 100 times (**Fig.1n,o**) to values of around 100 kΩ at 1kHz. These values were adjustable through modulation of electrodeposition parameters (Fig.1m) to match e.g. size constraints or input impedances of amplifiers. Surface modifications were mechanically stable during manipulation, tissue insertion, and even penetration in deep brain structures (∼1300 µm, **Supplementary figure 4**).

For extracellular recordings, microwires were assembled into low-profile custom 3D printed polymer holders incorporating a custom printed circuit board (PCB) and connector plug with matched channel count and a spacer separating individual fibers (**Fig.1e and Supplementary figure 1**). To record from distributed ensembles of neurons, we connected groups of 10-128 wires to custom designed PCBs using a modified ball-bonding technique (**Supplementary figure 7**). In this semi-automatic process, individual wires were placed in arbitrary 2D and 3D arrangements, at different depths with lateral spacing varying between 5 to 150 µm (shown for 50 µm inter-shank separation in **Fig.1e-i**). The overall layout of recording sites was tailored to the neuroanatomical structure of interest.

## Results

### Minimally invasive electrodes

To validate the technology for extracellular recording and stimulation, we inserted assembled jULIEs into the olfactory bulb (OB) of 4-6-week-old anaesthetized mice (**Fig.2a-g**). They reliably resolved single and multi-unit activity immediately after insertion starting from superficial layers (**Fig.2e**). This was in contrast to standard microfabricated probe recordings where unit activity is typically suppressed at up to ∼45 minutes after probe insertion^22^ which suggests that tissue integrity is better maintained upon jULIEs insertion.

**Figure 2.**
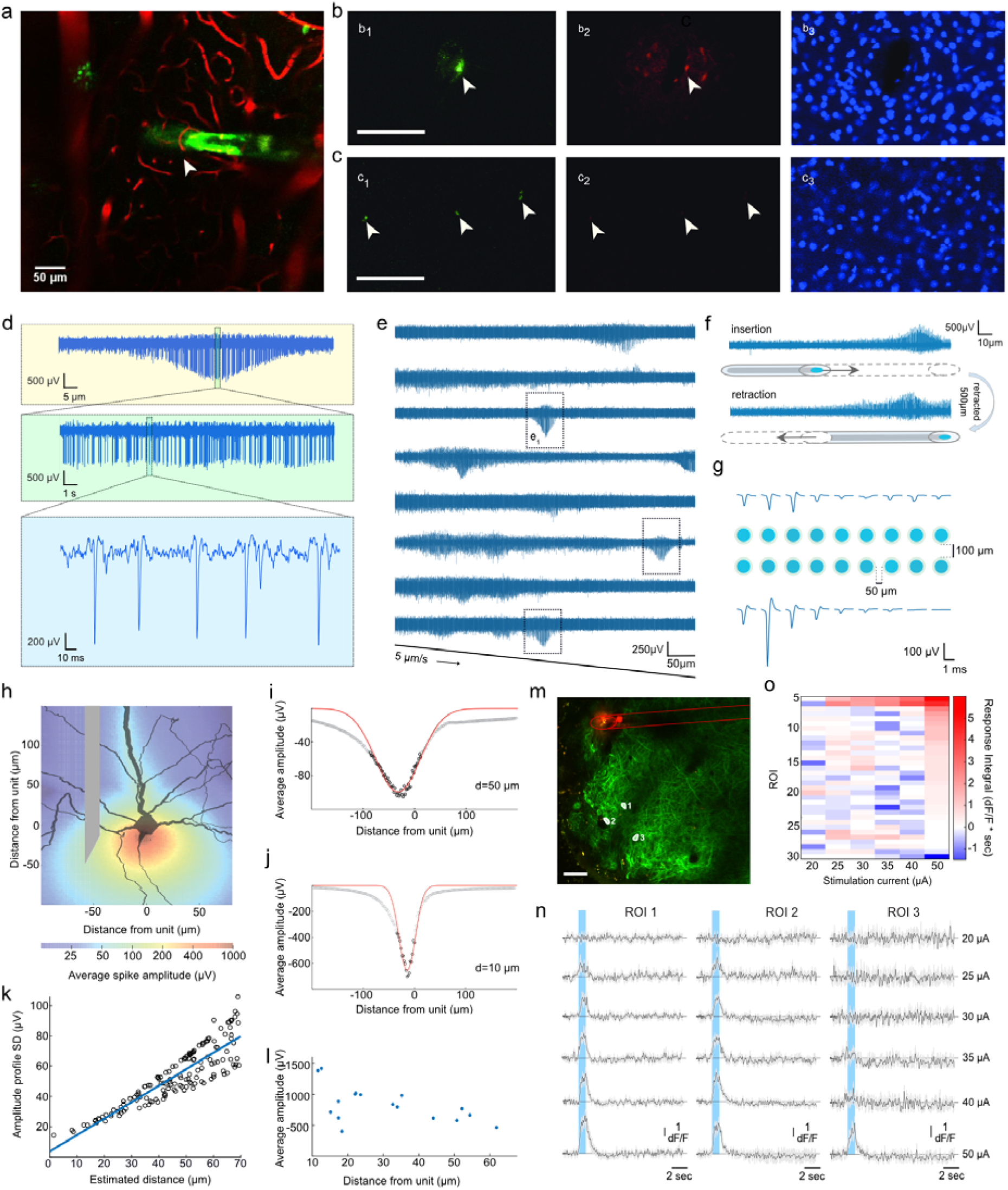
**(a)** *In-vivo* acute 2-photon microscopy observation of insertion of a DiO-labeled jULIE (green) into the MOB with sulforhodamine labeled capillaries (red). The jULIE was moved along its axis towards the middle of the OB then retracted several times (**see Supplementary Video 1**). **(b)** Acute tissue damage upon insertion of a Si-probe **(b**_**1**_**-b**_**3**_) and of a multi-shank jULIE probe (**c**_**1**_**-c**_**3**_; three jULIEs shown). Insertion sites are labelled by DiO (**b**_**1**_,**c**_**1**_; green). Damage to the blood brain barrier upon Si-probe insertion is indicated by the extravasation of albumin-bound EvansBlue (**b**_**2**_,**c**_**2**_; red), and displacement of cell nuclei (**b**_**3**_,**c**_**3**_; DAPI, blue). **(d)** Details of extracellular recording on a single channel during electrode insertion at constant speed ∼5 µm/s near a spontaneously firing unit. **(e)** Spontaneous activity on larger scale in the MOB recorded as multi and single-unit activity **(e**_**1**_) while jULIEs were inserted ∼400 µm into the OB. Spontaneously active units appeared as humps of increased amplitude on top of the multi-unit activity. **(f)** Spontaneous activity was preserved when jULIEs were inserted and retracted multiple times along the same axis in the vicinity of a spontaneously active unit; unit activity was recovered in comparable location during insertion and retraction (despite ∼500 µm further advancement of the jULIE). (g) Single-unit amplitude distribution on a custom arranged rectangular recording site layout featuring constant 50 µm lateral inter-shank and 100 µm inter-layer distance. **(h)** Heatmap of simulated spike peak amplitudes at different positions around a mitral cell. (**i,j)** Modelled spike amplitudes (black circles) at different positions along the electrode axis, and their Gaussian fit (red line) at 50 µm **(i)** and 10 µm **(j)** from the initial segment of biophysically realistic mitral cell. (k) Relationship of the standard deviation of fitted spike amplitudes profiles and the distance of the electrode axis to the initial segment of the neuron. **(l)** Peak amplitudes of maximal recorded action potentials from 17 jULIE probes and their estimated distance from the mitral cell initial segment, collected experimentally in the MOB. **(m-o)** Electrical stimulation in vivo in the OB using a single jULIE probe **(m)** In-vivo DiO labeled position of the jULIE stimulation site relative to ROIs labeled 1, 2, 3; **(n)** responses of the 3 ROIs indicated in **(m)** to increasing stimulation current; **(o)** and response of all identified cells contained within the field of view **(m)** to increasing stimulus current.

To directly assess the integrity of the vasculature, we coated jULIEs with DiO, labeled the blood vessels with sulforhodamine, and monitored tissue structure during insertion using 2-photon microscopy. While individual fibers were stiff enough to penetrate, they were sufficiently compliant to preserve both the structure of the tissue, with capillaries folding around the fiber shank, and slide on its surface without dragging the surrounding tissue, or apparent rupture of the vessels while inserted (**Fig.2a and Supplementary video 1**). To assess the latter more directly we performed histology after insertion experiments where the vasculature was loaded with Evans-Blue (**Fig.2b,c**), a dye unable to penetrate the blood-brain barrier due to its high-affinity binding to the large protein albumin^23^. For standard silicon probes (**Fig.2b**) insertion resulted in significant local damage to the blood-brain-barrier as measured by extravasation of Evans Blue (**Fig 2 b**_**2**_) and tissue displacement with loss of neuron density around the rectangular shank (**Fig 2 b**_**3**_). Insertion of the jULIEs, on the other hand (**Fig.2c**), left cell density seemingly unperturbed (**Fig.2c**_**3**_) and resulted in no detectable damage to the blood-brain barrier (**Fig.2c**_**2**_), suggesting that the neural tissue had been left structurally intact.

**Figure 3.**
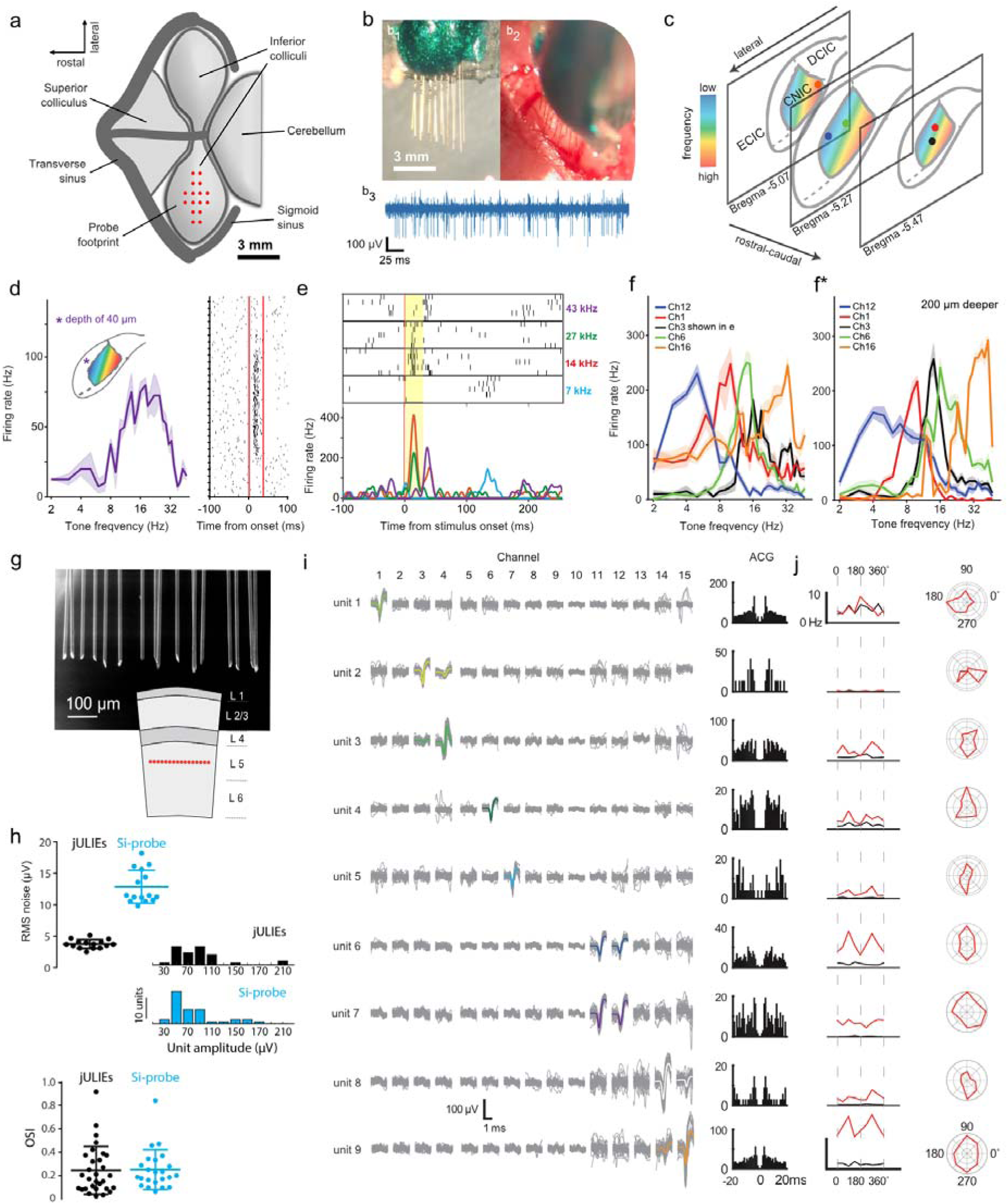
**(a-f)** jULIE recordings in inferior colliculus **(a)** Dorsal view of the inferior colliculus (IC) as seen after craniotomy. **(b1)**, Side image of the 16-channel jULIE probe; **(b2)**, top view of probe entering the dorsal surface of the IC; **(b3)**, voltage trace with units recorded from IC. **(c)** spatial distribution of recording sites used for the analysis in f and f*. Colors illustrate the arrangement of tonotopy in the IC, with low frequencies best represented in superficial / lateral regions and progressively higher frequencies in progressively deeper / more medial regions. The color of the dots in **(c)** corresponds to the color of the tuning curves shown in **(f)** and **(f*). (d)** jULIE recording from a superficial location (∼40 µm below dorsal surface; inset: schematic of recording location in IC dorsal cortex). Note the unimodal frequency tuning (left) and reliable, low-latency response (right); each dot represents a spike, red lines represent stimulus onset and offset respectively). **(e)** Evoked responses recorded from the same channel in the same location in response to 4 different tones (frequencies colored shown at the side of the raster plot; red line: stimulus onset; stimulus duration: yellow box), each repeated 5 times. Different frequencies evoked different response patterns. While 14 and 27 kHz elicited short-latency responses, 7 kHz elicited no response and 43 kHz elicited a late response. Sparse spontaneous activity is seen before and after the sound presentation. (f) Tuning curves (average of activity evoked by 5 sound presentations) recorded at different locations (color-coded in c) illustrating position-specific preferred frequencies. Tuning curves are pronounced, unimodal, and discriminable between units, despite being recorded simultaneously from several different positions in the inferior colliculus. **(f*)** Moving the jULIEs 200 µm deeper maintained receptive field properties. **(g-j)** jULIE recordings in visual cortex. (g) SEM micrograph of the recording site layout and their positioning in layer 5 (red dots in the bottom schematic). **(h)** jULIEs vs. Si-probe: RMS noise distribution, histogram of average spike amplitudes (negative deflection) from single units and distribution of orientation selectivity index (calculated from j) for all units recorded using jULIEs and Si-probe. **(i)** left: Waveforms of all units separated during a single jULIE penetration (650 µm depth). Average spike waveforms shown in color. Right: Auto-correlograms (ACG) of spike times for separated units. **(j)** left: Visual responses for the units shown in **(i)** for 8 different drifting directions (red) and baseline firing rate (black). Asterisks denote significant evoked activity in at least one direction. Right: Polar plots showing normalized tuning curves.

### In-vivo extracellular recordings

We recorded spontaneous single and multi-unit activity with high signal-to-noise ratio while slowly inserting jULIEs into the main olfactory bulb (MOB) (**Fig.2, Supplementary figure 5**). At insertion speeds of ∼5 µm/s we readily detected the emergence of unit activity (**Fig.2d,e**). Action potential (AP) amplitude increased from background levels to its maximum as jULIEs were advanced by 10-100 µm and subsequently decreased over similar length scales with further insertion. Highest amplitudes exceeded 1 mV suggesting close proximity of the recording site to the intact neuron at that point^24,25^.

When jULIEs were displaced from the surface towards the center of the olfactory bulb, we found that spiking activity emerged and disappeared in a symmetrical fashion, consistent with the recording electrodes passing through the volume without mechanically dragging the tissue, while capillaries folded and slid around the smooth shank (**Supplementary video 1**). When jULIEs were inserted and retracted multiple times on the same axis around a spontaneously firing unit, activity was preserved at the same location over several insertion-retraction cycles extending beyond the recorded units by several hundred microns, suggesting that the local neural network remained functionally intact, and the electrode neither caused significant damage to nor displaced neurons (**Fig.2f**). Using a jULIE array formed by two stacked layers of nine probes each with 50 µm inter-shank spacing and 100 µm inter-layer distance, single active units were resolved on several adjacent channels (**Fig.2g**). It is to be noted that such arrangements enabled the usage of well-established spike-sorting algorithms such as Klustakwik^26^, KiloSort^27^ or Plexon Offline Sorter (Plexon Inc.).

### Electrode - neuron distance estimation

To more precisely estimate the position of jULIEs in relation to the recorded neurons, we calculated the electrical field around biophysically realistic models of mitral cells **(Fig.2h)**, and estimated the resulting spike amplitudes during axial movement of the electrode **(Fig.2i,j).** Simulations of the extracellular electrical field demonstrated that the distance between the axis of electrode movement and the soma of the neuron could be predicted from the standard deviation of the amplitude profile **(Fig.2i-k).** We thus performed further electrode insertions into the olfactory bulb while monitoring the change of spike amplitude during motion, and estimated the distance from the recorded units **(Fig.2l).** We found that reliable recordings could be made at estimated (horizontal) distances as low as 10 µm from the axon initial segment. This suggests that sharpened, electrochemically modified microwires could indeed record in close proximity to intact neurons (hence juxtacellular Ultra-Low-Impedance Electrodes, jULIEs).

### Stimulation

A key advantage of the multi-step electrochemical modification described here is the large charge storage capacity due to the increased surface area and the stable reversibility of the Ir^3+^/Ir^4+^ redox couple. To assess whether this would indeed allow for electrical stimulation efficient enough to drive neuronal activity with ultramicroelectrodes, we performed systematic microstimulation under 2-photon observation in the olfactory bulb of transgenic mice expressing the genetically encoded Ca^2+^ indicator GCaMP6f in projection neurons (Tbet-Cre × Ai95). Using a single jULIE fiber **(Fig.2m)** 5-10× 1 ms pulses were injected carrying currents from 2 µA to 150 µA in 5 µA increments (see methods). Cells were found to respond to increasing stimulation levels with most cells in the proximity of the stimulation site responding at 50 µA injection current **(Fig.2m-o).** Electrical properties of the interface remained stable throughout 200,000 injected voltage pulses **(Supplementary figure 6a).** Moreover, electrochemical impedance spectroscopy and cyclic voltammetry characterization after current stimulation revealed only a threefold increase after 1 million pulses (Supplementary figures 6c,d), enabling the use of the same site for further recordings.

### Custom site layout

Silicon-probes are widely adopted because of their ability to perform extracellular recordings with relative ease on multiple channels. However, the underlying microfabrication methods have limitations in terms of materials, dimensions and required efforts to customize sensor layout to fit a certain target brain structure. In contrast, the layout of the recording sites of jULIEs can be readily customized **(Fig.1e-i)** to best match anatomical requirements of individual experiments. Microwires can be pre-arranged to maximize site lateral density using micron-scale polymer templates **(Supplementary figure 1)** which help splay fibers at insertion **(Supplementary video 2).** To record from sensory areas in the mouse brain, 16-channel pre-arranged probes were assembled: recording site layout was mapped onto the tonotopical arrangement in the auditory inferior colliculus (IC); or, as an evenly spaced one-layer arrangement, onto L5 neurons in the primary visual cortex.

### Recordings in the inferior colliculus

When sampling auditory physiology, it is often desirable to record along the tonotopic axis to resolve population activity evoked by different frequency components of an auditory stimulus. This often requires multi-shank probes which can be challenging in mice, as tissue damage around the multiple insertion sites is thought to limit the number of successfully recorded units^5,28,29^. Thus, to sample from the 3-dimensional volume of the IC, a 4-layer stacked jULIE probe with a rectangular footprint **(Fig.3a)** was assembled. Individual probe shanks **(Fig.3b)** were separated horizontally by ∼100 µm, recording sites were vertically arranged at 100 µm steps matching the slope of the isofrequency planes and inserted into the IC of anaesthetized mice with the deepest shank next to the medial axis, **(see Fig.3c).** Unit activity in the IC was readily resolved **(Fig. 3b3)** with presentation of pure tones. Notably, while recordings from superficial layers are generally difficult to obtain^30^, jULIEs allowed to record evoked spiking activity with clear tuning in superficial layers of the IC already at ∼40 µm from the surface (Fig.3d). Overall, the probe mapped onto the tonotopy of the IC (Fig.3e,f) allows to record tone evoked responses in units across the 2-dimensional surface at different depths (Fig.3,f and f*), including at very superficial layers.

### Recordings in the visual cortex

The neocortex is a layered structure where targeting individual layers with many recording sites for extracellular recordings is challenging, yet often desirable. While e.g. Si-polytrodes^31^ can achieve single layer multi-channel recordings in cortex, their larger footprint results in local tissue damage and limits their versatility. jULIEs in turn can be arranged in arbitrary 2D and 3D patterns, such as a single-layer horizontal site layout **(Fig.3g)**, to target recording sites to individual layers. Recording sites can be spaced as close as necessary, not limited by the number of shanks.

We tested the utility of the laterally dense probes by recording from the primary visual cortex (V1) of lightly anaesthetized mice at cortical depths corresponding to layer 5 (500-750 µm). Following surgery to expose the cortex, a jULIE probe (16 wires arranged horizontally with 50 µm inter-shank spacing) was lowered into the cortex. Separate recordings were performed under similar conditions with a comparable 16-channel Si-probe (4 shanks 150 µm apart, each containing 4 recording sites in a tetrode arrangement, see Online Methods). Several recording sessions, at least 50 µm apart in depth, were made both with the jULIEs and the Si-probe.

Recordings from individual jULIEs had lower noise (RMS 3.77 ± 0.69 µV; n = 15 wires) compared to the Si-probe used in this experiment (RMS 12.88 ± 2.64 µV; n = 15 sites, Fig.3h). Single units appeared on multiple wires (e.g. unit7, Fig.3i), and the same wire could record from multiple single units (e.g. ch15, **Fig.3i**). Average spike amplitudes of well-separated units showed a comparable distribution to those from the Si-probe (**Fig.3h**, jULIEs: 66.39 ± 45.48 µV, n = 33; Si-probe: 77.44 ± 40.40 µV, n = 39; *p* = 0.28, Student’s t-test).

Well-isolated single units (see Online Methods for sorting) could be detected on single or multiple (usually 2) neighboring recording sites **(Fig.3i).** We recorded visually evoked single-unit activity using standard drifting gratings (8 directions, see Online Methods for details). As expected^32^, activity evoked by visual stimuli showed characteristic tuning curves (**Fig.3j** right) for single units. Units recorded on the same channel (e.g. units 6 and 7) had different levels of evoked activity and markedly different tuning, consistent with the assumption that units represent individual neurons. Furthermore, drifting-grating responsive single units recorded from jULIEs and the Si-probe had similar distribution of orientation selectivity index (jULIEs OSI: 0.24 ± 0.20, n=30 cells; Si-probe OSI: 0.25 ± 0.17, n=23 cells; *p* = 0.88, Student’s t-test, **Fig. 3h**).

## Discussion

Here we have described the manufacturing, modification and use of “jULIEs”, glass-metal composite ultramicroelectrodes, as a minimally invasive and scalable approach to record high-quality extracellular signals from the intact mouse brain. We demonstrated that electrochemical multi-step modifications with nanomaterials substantially reduced interfacial impedances, decoupling wire dimensions and materials from electrode-electrolyte coupling. jULIEs were compliant with the tissue structure, causing no detectable tissue displacement or damage to the blood-brain-barrier, making them an ideal candidate for long-term implantation. This was likely due to the overall small dimension of individual fibers^29,33^, cylindrical form, sharpened tip and smooth shanks which have been suggested to keep mechanical interaction with surrounding tissue minimal^10^. Other advantages of modified microwires are their composite nature: metal cores are effectively fused with the glass ensheathing providing a robust, defect and delamination-free very low parasitic-capacitance insulation^34^. Polymer coated microwires^35^, microfabricated polymer electrodes^36-38^, carbon fibers^33,39^, multimodal pipettes^40^, syringe-injectable mesh electrodes^41,42^ and niobium microwires^43^, amorphous SiC^44^ and active-CMOS probes^45^ are promising avenues for potentially low-damage neural recordings. Several approaches have been put forward to allow for their connectorization to standard readout electronics however at very large scale (above 10k) these still remain a challenge. Due to their metal core, the jULIEs can be readily connected to integrated read-out electronics in a scalable manner, either through direct bonding on a smaller scale **(Supplementary Fig 7)** or through non-thermal flip-chip bonding at very large scale^11,34^. Moreover, when assembled into customized neural probes, the overall layout of sensor sites can be individually tailored to fit the neuroanatomical structure in three dimensions.

Multi-step electrochemistry enables decoupling of the recording interface in contact with the neuronal tissue from the material properties of the wires and their limitations. This might further allow usage of core compositions which improve wetting capabilities needed to manufacture thinner fibers^46^. NanoAu deposition could then restore the contact surface area and electrochemical compatibility, and, in tandem with IrOx deposition, provide a low impedance electrode-electrolyte interface despite submicron core diameters. In fact, microwire modification with an inert metal such as the nanoAu enables access to a diverse set of impedance reducing strategies with materials such as polymers, carbonaceous or hybrid materials. Furthermore, inert metal cores have very low (<100 Ω/m) axial resistance and are compatible with a wide range of further possible biofunctionalization strategies, to detect e.g. pH^47^, catecholamines^48^, or DNA^49^. Finally, through multi-step electrochemistry we created high rugosity surfaces in combination with a reversible redox couple at the electrode-tissue interface, resulting in an attractive and stable substrate for microstimulation purposes.

Low invasiveness allowed us to record single units on several adjacent jULIEs, despite the multiple penetrations **(Fig.2g)**. In combination with the flexibility of geometric custom arrangement of recording sites, this could be used to tailor probe arrays for optimal unit isolation, maximizing unit yield^50^.

For understanding the brain, recording and stimulation in three dimensions at high speed, and at a large distributed scale is critical. Various imaging, electrical or genetic approaches are available, each challenged by particular limitations of scalability^50^. Currently, the technologies that not only might be closest to that goal but are also applicable to clinical challenges are electrical recording and stimulation techniques with invasiveness considered to be the central obstacle. Here, jULIE probes combine low invasiveness with high quality electrical recording and stimulation, thus offering a platform for interfacing with brain activity at brain-wide scale with single-cell resolution.

## Supporting information

jULIEs insertion in-vivo under 2p

## Acknowledgements

We thank Martyn Stopps for help with electronic design, Isabell Whiteley for technical help with histological processing, and Anatolii Ioisher for advice on glass-metal microwires. We also thank Lucy Collison and the EM STP team for help with SEM imaging, the Making Lab and the BRF for technical help. We also thank to Ecaterina Ware and Mahmoud Ardakani from Imperial College Faculty of Engineering, Department of Materials for their help with characterizing and imaging nanoAu and IrOx nanostructures. We thank Howard Marriage and Veronique Birault and the Translational team at the Crick for supporting development of jULIEs through the *Idea 2 Innovation* grant series. This work was also supported by the Francis Crick Institute which receives its core funding from Cancer Research UK (FC001153), the UK Medical Research Council (FC001153), an HFSP grant (RGP 00048/2013), an NIH BRAIN initiative grant (1U01NS094248-01) and the Wellcome Trust (FC001153) and the Medical Research Council (MC_UP_1202/5). Ede A. Rancz is a Sir Henry Dale Fellow (Wellcome, 104285/B/14/Z). Andreas Schaefer is a Wellcome Trust investigator (110174/Z).

## Dissemination

jULIE probes in standard and custom layouts can be obtained from neurotrodics.com.

## Materials and methods

### Animal welfare

All experiments were performed according to United Kingdom Home Office regulations (Animal, Scientific Procedures Act 1986) or the guidelines of the German animal welfare law and were approved by the local welfare committees and veterinarians.

### jULIE neural probes

Glass-metal composite ultramicroelectrodes were fabricated by an adapted Taylor-Ulitovsky thermal drawing method^8^. Typically, a cylindrical borosilicate glass tube (OD 10 mm, ID 6 mm, Pyrex, Corning, UK) was loaded with a metal rod (Puratronic, Alfa Aesar, UK) and inductively heated (PowerCube 900, CEIA, UK) to 850 – 1000 °C until a separating drop formed. The thermoformed part was removed and the flowing fiber was, at high speeds (2 km/min), continuously spooled up on a drum as depicted in **Figure 1a.** This process resulted in electrically continuous conductive ultramicroelectrodes with outer diameters ranging from 10 µm to 100 µm with cores from 2 µm to 10 µm and lengths in the order of hundreds of meters.

Ultramicroelectrodes were bundled together using an Optima 1100 (Synthesis, India) winding machine and embedded in Crystalbond 509 (Agar, USA) a thermoplastic, dissolvable resin to fit custom 3D-printed polishing holders. jULIEs were sharpened at 30 degrees using a MetaServ 250 polisher equipped with a Vector head (Buehler, USA) and sequentially polished at 5 different levels using particles (25 µm, 9 µm, 3 µm, 1 µm, 0.05 µm) suspended in water-based emulsion (Buehler, USA) for 40 seconds each. Sharpened wires were de-embedded by solubilization (1000:1 solid/liquid ratio) in Crystalbond 509 organic stripper (Agar, USA) for 24 h then washed with isopropanol and dried at 60 °C overnight before long-term storage.

To record *in vivo* extracellular signals, we assembled jULIEs (**Figure 1e-i, Supplementary Figure 1)** into neural probe modules of 16, 32 and 128 channels onto custom PCB (E44, LPKF Protomat, UK) using a modified instrument and bonding procedure (F&S Bondtec, UK) which allowed in-situ read-out of connectivity and was equipped to remove glass insulation. Bonded and sharpened wires were then modified electrochemically as described below.

### NanoAu electrodeposition

NanoAu was electrodeposited from a two-part aqueous cyanide bath containing 50 gL^-1^ potassium dicyanoaureate(I) (K_2_[Au(CN)_2_]) and 500 gL^-1^ KH_2_PO_4_ dissolved sequentially in ultrapure deionized water (18 MΩ·cm) (Tech, UK) at 60 °C. All reagents were supplied by Sigma-Aldrich, UK and used without further purification. Prior to electrodeposition the polished substrate was washed with deionized water, rinsed with ethanol (90%), wiped with a lint-free cloth (Kimwipes, Kimtech, UK) and dried at 50 °C for 1 hour in an oven (Memmert, Germany). The electrodeposition protocol was carried out with a multichannel potentiostat-galvanostat (VSP 300, Bio-Logic, France) controlled with EC-Lab software (Bio-Logic, France) in a three-electrode cell setup composed of the assembled gold jULIE probes as working electrodes (W_E_), a coiled 1 mm thick platinum wire (PT005150, 99.95%, Goodfellow, US) as counter electrode (C_E_) and a Ag/AgCl|KCl_3.5M_ reference electrode (REF) supplied by BASi, USA (E vs. Normal Hydrogen Electrode (NHE) = 0.205V). The REF was kept separated from the bath by a glass tube containing the support electrolyte and a porous Vycor glass separator. For nanoAu deposition the W_E_ potential was kept at E_red_ =-1.1 vs. REF for up to 180 seconds according to the desired hemisphere size as shown in **Figure 1m**. The electrodeposition bath was maintained at 60 °C using a thermostat under vigorous (500 rpm) stirring.

### IrOx electrodeposition

The electrodeposition protocol was carried out from a modified electrolyte solution based on a formulation reported by (Kazusuke 1989; Meyer 2001) containing 10 gL^-1^ iridium (IV) chloride hydrate (99.9%, trace metal basis, Sigma-Aldrich, Germany), 25.3 gL^-1^ oxalic acid dihydrate (reagent grade, Sigma-Aldrich, Germany), 13.32 gL^-1^ potassium carbonate (99.0%, BioXtra, Sigma-Aldrich, Germany). Reagents were added sequentially to 50% of the solvent’s volume firstly by dissolving IrCl_4_ in oxalic acid followed by the addition of K_2_CO_3_ over a 16-hour period until pH=12 was reached. The electrolyte was aged approx. 20 days at room temperature in normal light conditions until the solution reached dark blue color. IrOx was electrodeposited using the instrumentation and setup described above (see section nanoAu Electrodeposition). The deposition protocol was composed of two consecutive stages combining cyclic voltammetry (CV) and a pulsed potentiostatic protocol (PP). Between protocols, the W_E_ was kept at the open circuit voltage (OCV) for 180 seconds to allow equilibration. During CV deposition the W_E_ potential was cycled 50 times between -0.5 V and 0.60 V vs. REF at 1 Vs^-1^ in both anodic and cathodic directions. During the pulsed potentiostatic deposition the W_E_ potential was stepped 250 times between 0 V to 0.60 V vs. REF in 1 second steps.

### jULIEs electrochemical characterization

Electrochemical behavior of the sensor sites was monitored individually in the unmodified state (polished gold/glass surface), after modification with nanoAu, and after modification with nanoAu+IrOx. Multichannel jULIE modules were assembled and connected to the potentiostat using a custom PCB and matching connector plugs. Characterization was done in 150 mM phosphate buffered saline using a 3-electrode cell-setup (see details for C_E_ and REF above) for their cyclic voltammetry response and electrochemical impedance (EIS) profile. EIS measurements were performed by applying a 10 mV sine-wave around the open circuit voltage (OCV) in the frequency range 1 Hz to 100 kHz with average 3 measurements per frequency and 5 repetitions for each channel. Using an identical cell setup, cyclic voltammetry response was recorded for individual channels by sweeping electrode potential from -0.5 V and 0.6 V vs. REF with 100 mVs^-1^ sweeping rate in both cathodic and anodic directions.

### FIB-SEM and STEM characterization

Modified sensor sites were characterized for their microstructure, atomic lattices and chemical composition by field emission electron microscopy (FESEM) and scanning transmission electron microscopy (STEM) using a multipurpose 200 kV JEOL JEM-2100F TEM analytical electron microscope coupled with an Energy Dispersive X-ray Spectrometer (EDS) and Oxford Instruments INCA/Aztech EDS 80 mm X-Max detector system for elemental analysis with nanometer spatial resolution. A dual-beam FEI Helios 600 FIB/SEM system equipped with a gallium ion source operating in the accelerating voltage range 0.5–30 kV and an Omniprobe™ micromanipulator was used to morphologically characterize and prepare samples for transmission electron microscopy (TEM) imaging. Sample preparation consisted of: (i) deposition of a protective platinum (Pt) layer by sputtering on to the specimens, (ii) milling a thin slice perpendicularly to the sample surface, (iii) extraction and gluing the specimen slice to a TEM grid, and (iv) further thinning of the sample with low-voltage focused ion-beams at grazing incidence until an electron-transparent region was obtained. These steps were repeated for both nanoAu and nanoAu+IrOx specimens as depicted in **Supplementary figures 2 and 3.**

### LFP modelling

For simulation of the LFP around a detailed mitral cell model, we used a Neurolucida reconstruction of a mitral cell **(IF04208**^**51**^). Ion channel densities for different domains (glomerular tuft, apical dendrite, lateral dendrite, soma, axon) were adapted from Rubin et al. 2006. To gain a more accurate picture of the field around the initial segment, a sodium channel density of 2000 pS/µm^2 52,53^ was included in the first 5 µm of the axon. The LFP was simulated at different locations with the line-source method^54^, with the LFPy Python package^55^.

### Histology

Tissue integrity post jULIE insertion was determined by histological methods as described^34^. In brief, after craniotomy, 0.2ml of 0.5% Evans Blue was injected into the tail vein. Prior to insertion, jULIEs were dipped into SP-DiO (Molecular Probes, OR, USA) and allowed to dry. After the jULIEs were removed from the brain the mouse was perfused with ice cold 4% PFA, the brain was harvested and stored in 4% PFA overnight. Using a Vibratome (Leica, Germany), the brain was sliced into 100 µm horizontal sections. Slices were stained with DAPI using a 1:1000 DAPI:PBS wash for 10 minutes, transferred to fresh PBS, mounted and sealed. Imaging was completed on a confocal microscope (Leica SP5).

### Extracellular recordings in the olfactory bulb

To test jULIEs we performed recordings *in vivo* in the OB of mice. 4-6 weeks old mice were anaesthetized using a mixture of Ketamine/Xylazine (100 mg per kg of body weight) and xylazine (20 mg per kg for induction and 10 mg per kg for maintenance) administered intraperitoneally and supplemented as required. Body temperature was maintained at 37°C using a feedback-regulated heating pad (FST, USA). The main olfactory bulb was accessed through a 2×2 mm craniotomy window after fixation of the skull with a head-plate. The surface of the brain was protected by an imaging well containing Ringer’s solution and caudally fixed Ag|AgCl reference electrode.

Using micromanipulators (Luigs & Neumann, Germany) the jULIE probes were lowered to a depth of approximately 400 µm. Extracellular recordings were performed using a Tucker Davis RZ2 amplifier with a RA16AC-Z headstage or an Intan RHD2132 headstage on an OpenEphys amplifier (openephys.org).

Mice were presented with mixtures of odorants (Mixture 1: Ethyl butyrate & 2-hexanone, Mixture 2: Eucalyptol & Amayl accotate) (Sigma Aldrich, USA) diluted 1:5 with mineral oil. Recordings were taken from 400 µm from the surface of the bulb. Units were isolated using either Spike2 (Cambridge Electronics Devices, Cambridge, UK) or Kilosort (Pachitariu 2016) and units with well-defined auto-correlograms were selected for further analysis. Units were found with distinct responses to different odors, that were stable across repeats.

### Extracellular recordings in the inferior colliculus

Female C57BL/6JOlaHsd (Janvier, France) mice were anaesthetized with Avertin (0.15 ml/10 g). Additional doses (0.03 ml/10 g) were given as needed to maintain anesthesia. After anesthesia, the animal was fixed with blunt ear bars on a stereotaxic apparatus (Kopf, Germany). Body temperature was maintained at 36°C with a feedback-regulated heating pad (ATC 1000, WPI, Germany). Vidisic eye gel (Bausch + Lomb GmbH, Germany) was used to prevent the eyes from drying out. A metal head-holder was glued to the skull 1.0 mm rostral to Lambda with methyl methacrylate resin (Unifast TRAD, GC). A craniotomy of 0.8 mm × 1.0 mm with the center 0.85 mm from the midline and 0.75 caudal to Lambda was made to expose the left inferior colliculus. Dura was carefully removed and the surface of the brain protected with Saline (B. Braun, Germany). The inferior colliculus was identified by its position posterior to the transverse sinus and anterior to the sigmoid sinus. With a micromanipulator (Kopf, Germany), jULIEs were lowered vertically and advanced into the inferior colliculus.

The electrophysiological signal was amplified (HS-18-MM, Neuralynx, USA), sent to an acquisition board (Digital Lynx 4SX, Neuralynx, USA), and recorded with a Cheetah 32 Channel System (Neuralynx, USA). The voltage traces were acquired at a 32 kHz sampling rate with a wide band-pass filter (0.1 ± 9,000 Hz).

The sound was synthesized using MATLAB, produced by a USB interface (Octa capture, Roland, USA), amplified (Portable Ultrasonic Power Amplifier, Avisoft, Germany), and played with a free-field ultrasonic speaker (Ultrasonic Dynamic Speaker Vifa, Avisoft, Germany). The speaker was positioned 15 cm away from the right ear. Sound intensity was calibrated with a Bruël & Kjaer microphone. For measuring the tonal receptive field, we used sound stimuli consisting of 30 ms pure tone pips with 5 ms rise/fall slope repeated at a rate of 2 Hz. Thirty-two frequencies (2 kHz to 47 kHz, 0.16 octave spacing) were played in a pseudorandom order at intensities of 60 or 70 dB. Each tone-intensity combination was played 5 times.

The recorded voltage signals were high-pass filtered at 500 Hz. The root-mean-square (RMS) of the noise of each channel was calculated as the RMS level of the filtered trace during the first 10 seconds of recordings, which included spontaneous and evoked activity. To improve the signal-to-noise ratio of the recording, the common average reference was calculated from all the functional channels and subtracted from each channel (Ludwig 2009). For multiunit analysis, spikes were detected as local minima below a threshold of 6 times the median absolute deviation of each channel. If the calculated value was higher than -40 µV, the threshold was set to -40 µV. To analyze the sound-driven responses, peri-stimulus time histograms (PSTHs) were built by aligning the signals at stimulus onset and calculating the number of spikes/ms. Iso-intensity tuning curves were built from the sum of spikes in an 80 ms window from stimulus onset, at each intensity as a function of frequency.

### Extracellular recordings in the L5 of primary visual cortex

Male C57BL/6 mice between 2-4 months old were used for acute recordings in the visual cortex. For surgery, mice were anaesthetized with isoflurane (3% induction followed by 2% maintenance), and fixed on a stereotaxic apparatus (Model 940, David Kopf instruments, Germany) using ear bars. Body temperature was maintained at 36 °C with a DC temperature regulation system (FHC, Inc. USA). A skin incision was made and the exposed skull was cleaned and dried. A metal headplate was cemented to the skull using dental cement (Super-Bond C&B, Sun Medical, Japan). Animals were allowed to recover after surgery in their home cage. Analgesia (2 mg/kg Meloxicam with 0.1 mg/kg Buprenorphine) was provided. On the day of the recording, a craniotomy 1.5 mm long, 0.3-0.5 mm wide was made with small drill bits on the right hemisphere under isoflurane anesthesia (3% induction followed by 2% maintenance). The long axis of the craniotomy was parallel with the lambda suture. The dura was removed and the craniotomy covered with a silicone-based sealant (Kwik-cast, World Precision Instruments). Following recovery from surgery (4-24 hours), the animal was lightly anaesthetized in 1.0-1.5% isoflurane and head-fixed to the recording setup. The craniotomy was kept moist with cortex buffer (NaCl 125 mM, KCl 5 mM, HEPES 10 mM, MgSO_4_ 2 mM, CaCl_2_ 2 mM, Glucose 10 mM, pH 7.4) and the jULIEs or Si-probe (A4X1-tet-3mm-150-121, Neuronexus, USA) was slowly (∼10-20 µm/minute) inserted into primary visual cortex using a micromanipulator (SM1, Luigs & Neumann, Germany). Neural signals were recorded using a PZ2-32 preamplifier and an RZ2 BioAmp Processor (Tucker-Davis Technologies, USA). Data were acquired at 24.4 kHz sampling rate and recorded using TDT’s OpenEx software suite. The amplifier ground was connected to a screw implanted in the skull.

For visual stimulation full screen drifting gratings of 8 different directions (spatial frequency: 0.08 cpd; temporal frequency 2Hz; 2s long, with 2s grey screen inbetween), were presented in randomized order on a 27-inch LCD screen (E2711T, LG Electronics), placed 15-25 cm from the left eye. Gratings were generated and presented using MATLAB (Mathworks, USA) and the Psychophysics toolbox^56^.

Data were extracted and processed using custom written Matlab scripts.Neural signals were bandpass filtered at 300-5000 Hz for spike detection. Semi-automatic spike sorting was carried out using Klusta software with default parameters (Rossant 2016). Clusters with clear refractory period in the auto-correlogram (ISI violation <0.5%) and isolation distance >20 were classified as single units. Manual curation of resultant clusters was done using Phy^26^.

### In vivo two-photon olfactory bulb wire insertion and imaging

Mice were anaesthetized using a mixture of Fentanyl/Midazolam/Medetomidine (0.05 mg/kg / 5 mg/kg / 0.5 mg/kg). The skull overlying the dorsal olfactory bulb was thinned using a dental drill and removed with forceps, the dura was peeled back using fine forceps. Body temperature was maintained at 37°C throughout the experiment using a feedback-controlled heating pad (FHC, Inc. USA). Sulforhodamine 101 (Sigma Aldrich, 100 µm final concentration) was injected intraperitoneally to label blood vessels. Animals were then moved to a two-photon microscope (Scientifica Multiphoton VivoScope) coupled with a MaiTai Deep See laser (Spectra Physics, Santa Clara, CA) tuned to 940 nm (∼50 mW average power on the sample) for imaging. Images (512 x 512 pixels) were acquired with a resonant scanner at a frame rate of 30 Hz using a 16x 0.8 NA water-immersion objective (Nikon). For in vivo z-stack imaging, images were taken at a resolution of 512 x 512 pixels with 2 µm z intervals. Wires were dip-coated in DiO (Sigma Aldrich) before insertion into the dorsal olfactory bulb using a micro manipulator (Scientifica, Uckfield, UK). Images were analyzed post hoc using ImageJ (NIH, Bethesda).

### In vivo two-photon imaging of stimulation in the olfactory bulb

Mice were prepared for imaging as described in the previous section. Wires were dip-coated in DiI (Sigma Aldrich) before insertion into the dorsal olfactory bulb using a micro manipulator (Scientifica, Uckfield, UK). A silver/silver chloride counter electrode was inserted into caudal parts of the imaging well. For electrical stimulation, typically 100 ms long step current pulses were applied repeatedly to the inserted microwire relative to the caudal reference electrode with a Digitimer NL800A stimulator while imaging at typically 6 z-planes (∂z = 25 µm). Total frame rate was 30 Hz, resulting in an effective volume repetition rate of 5 Hz. Stimulation strength was varied between 5 and 85 µA in 5 µA steps and repeated 3-5 times for each stimulation strength. ROIs were selected manually offline using custom written routines in ImageJ and data was exported for further analysis in Matlab.

