## Supplementary materials for "jULIEs: extracellular probes for recordings and stimulation in the structurally and functionally intact mouse brain"

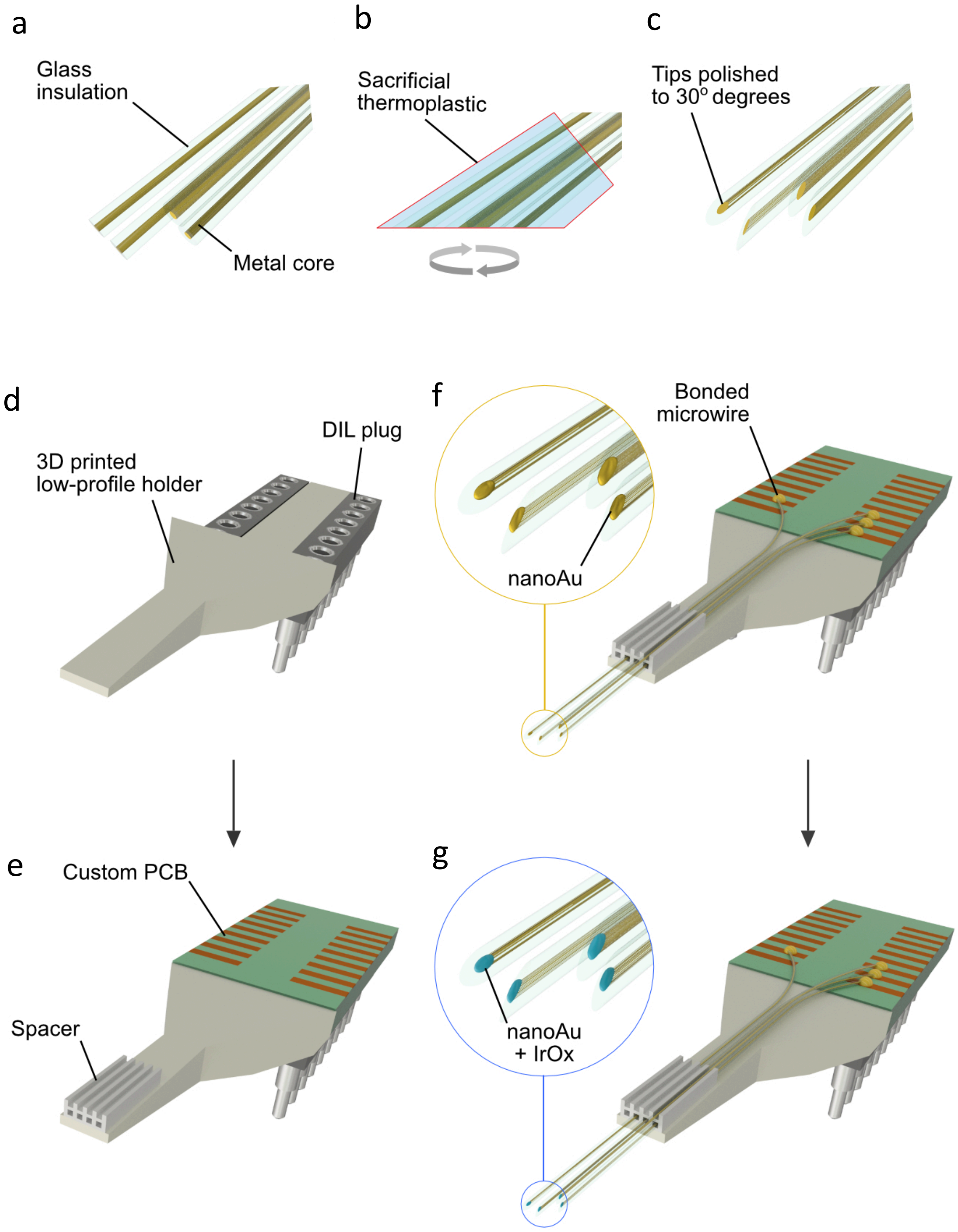


**Supplementary Figure 1**. Preparation and assembly of jULIE probes. **(a)** Microwires were grouped into bundles containing several thousand wires and embedded into Crystalbond 509. **(b)** Samples were fixed into a holder and polished on a Buehler 250 using 5 grits. **(c)** Sharpened wires were de-embedded overnight by solubilization in Crystalbond Stripper. **(d,e)** Low profile microwire holders, a custom PCB and connector plug were assembled. **(e)** Wire spacers were placed at the tip of the 3D printed holder. **(f)** Microwires were inserted into spacers and built up layer by layer. **(g)** Microwires were bonded and modified with nanoAu and IrOx and characterizes by EIS.

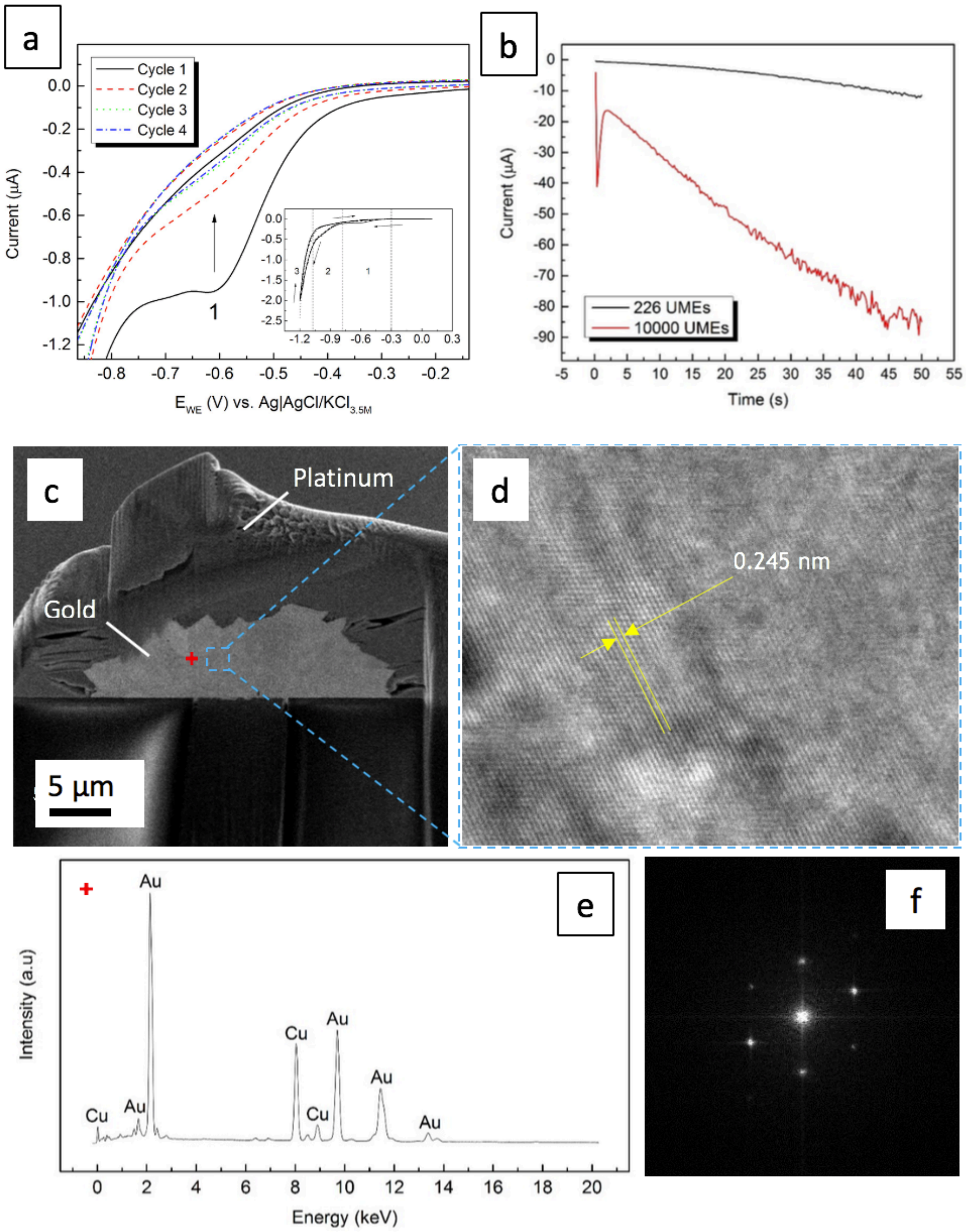


**Supplementary Figure 2.** Two-step nanoAu electrodeposition on polished gold/glass ultramicroelectrodes from an additive free cyanide electrolyte **(a)** Peak current (1) response for nanoAu electrodeposition during first voltammetry cycle at 50mVs$^{-1}$ between -1.2V and 0.2V vs. REF. **(b)** Potentiostatic plating of nanoAu on bundles containing several tens of individual glass-gold jULIEs c) FIB-SEM processed electron-transparent slice from the nanoAu deposit. Note the protective Pt layer deposited to facilitate FIB milling **(d)** STEM image of Au atomic lattice arrangement **(e)** EDS spectra of point marked by a red cross in (c), **(f)** FFT X-ray diffraction pattern indicating the single crystal nature of sample from panel (**d**)

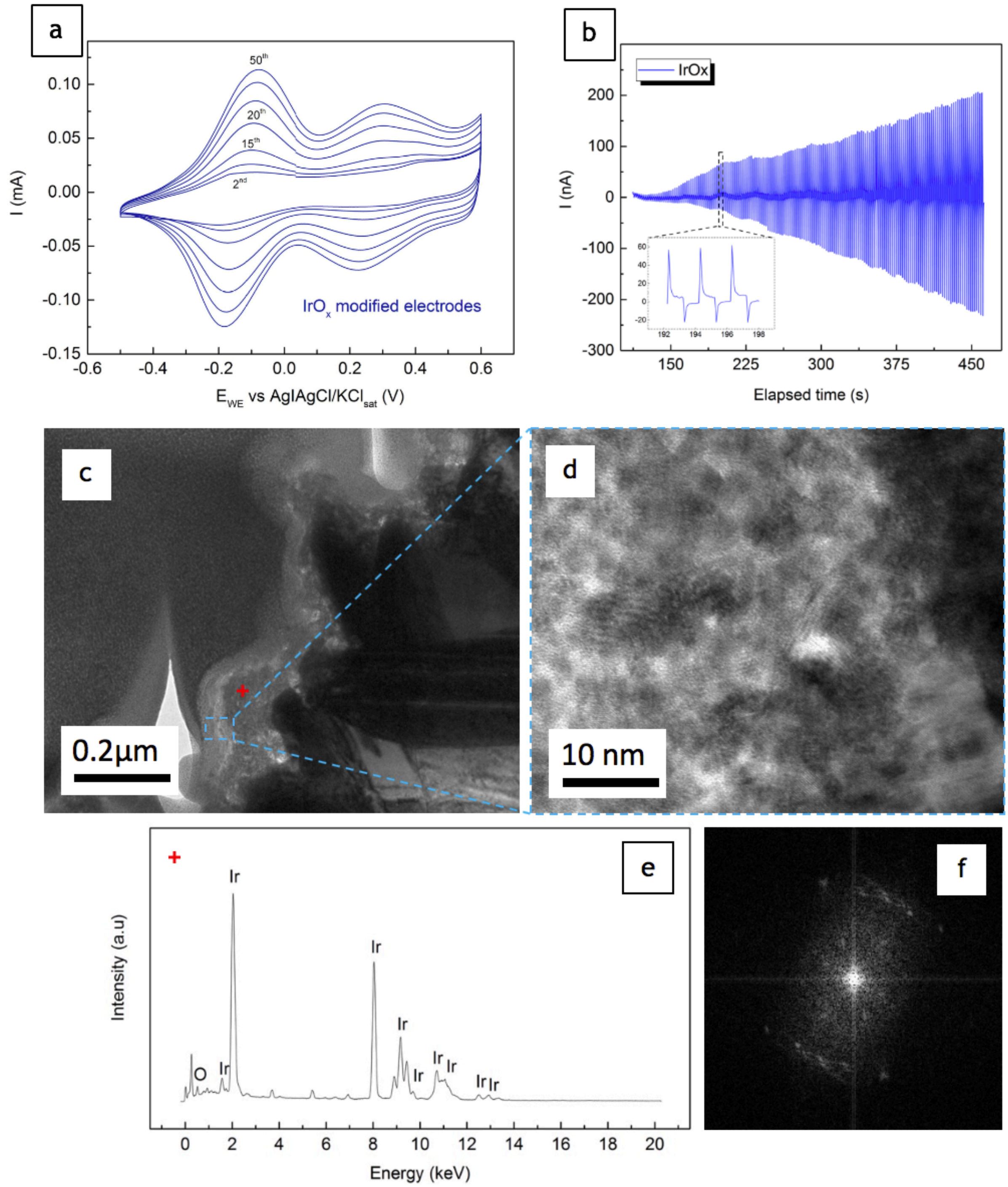


**Supplementary figure 3.** Two-step IrOx electrodeposition on nanoAu **(a)** Cyclic voltammetry electroplating on nanoAu at a sweep rate of $50 mVs^{-1}$ between -0.5V and 0.6V vs. REF; **(b)** Potentiostatic IrOx pulse-plating on nanoAu; **(c)** STEM micrograph of the electrodeposited $IrO_x$ thin-film; **(d)** STEM of bulk $IrO_x$ thin-film showing polycrystalline aggregates irregularly packed resulting in the porous overall structure; **(e)** energy dispersive x-ray diffraction spectra of the $IrO_x$ thin-film at position indicated by a red cross in (c); **(f)** FFT X-ray diffraction of polycrystalline $IrO_x$ showing typical polycrystalline response pattern.

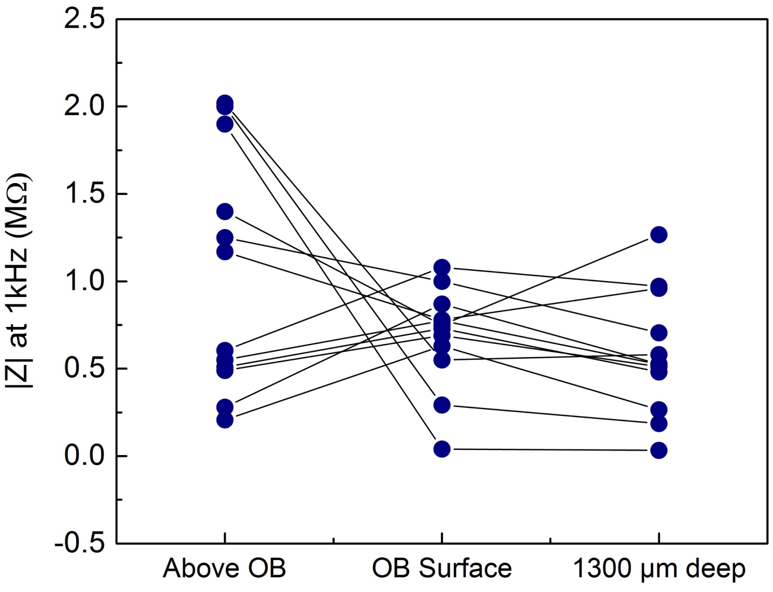


**Supplementary figure 4.** Impedance stability of the jULIE probes at different stages during an *in-vivo* experiment in the mouse olfactory bulb: before insertion in ACSF ("Above OB"), touching the surface of the brain (dura removed) and after insertion and axial displacement of ~1300 µm into the olfactory bulb.

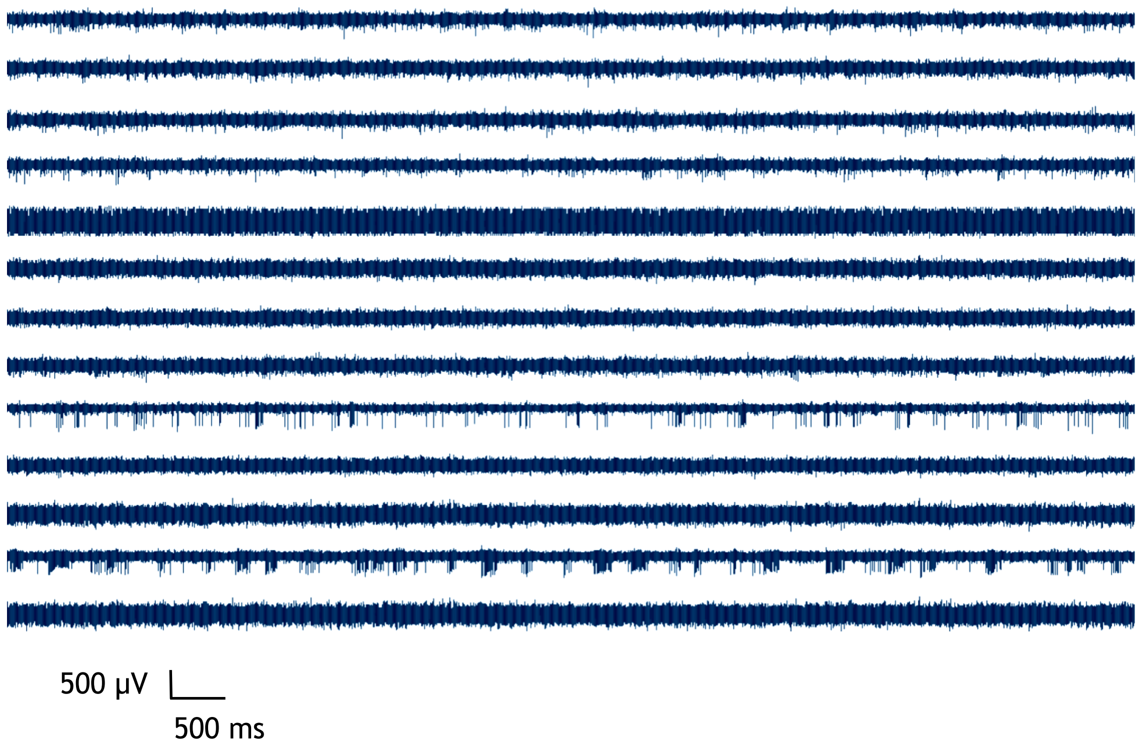


**Supplementary figure 5.** *In-vivo* extracellular recordings at ~400 µm deep in the main olfactory bulb of an anesthetized C57BL/6 mouse using a 13 channel jULIE.

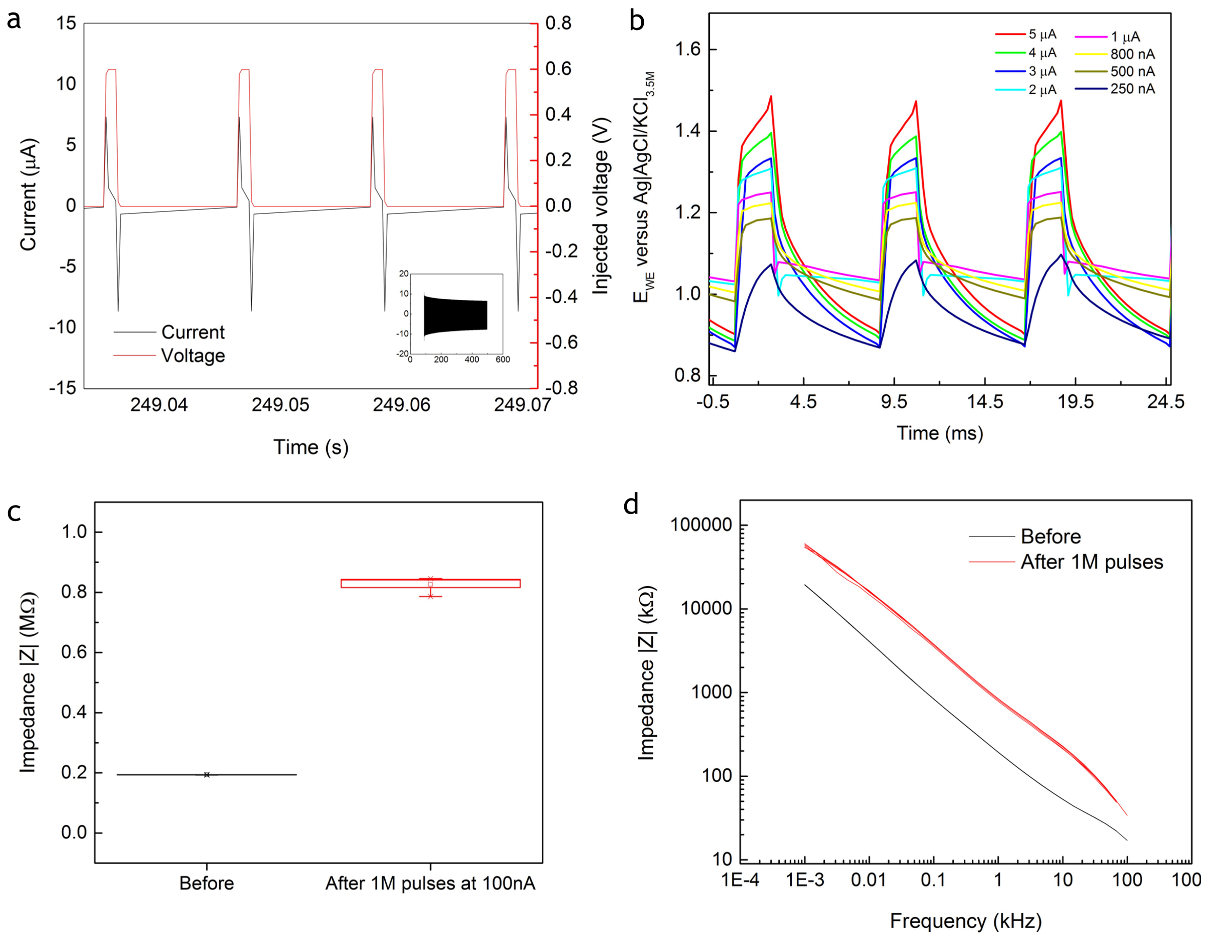


**Supplementary figure 6.** **(a)** Characterization of voltage stimulation performance of nanoAu + IrOx in cortex buffer for injected 1 ms long 600 mV square pulses. **(b)** Electrode voltage response to different injected currents. **(c)** Impedance response after 1M 100nA injected pulses and overall EIS response **(d)**.

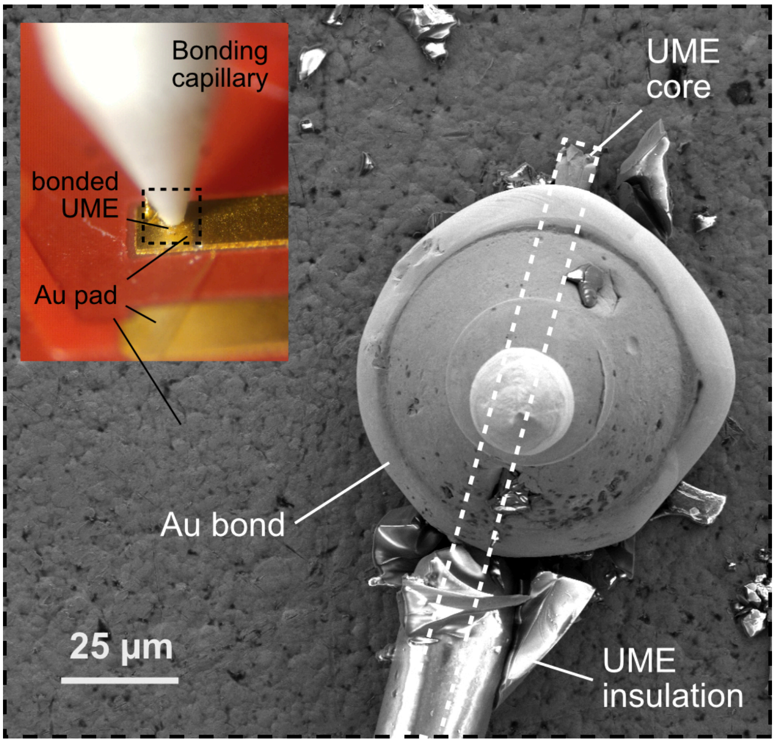


**Supplementary figure 7.** jULIE wire core bonded to gold pad after removal of glass insulation. Dry contact resistances range between 1Ω and 8Ω
